# Multiphoton minimal inertia scanning for fast acquisition of neural activity signals

**DOI:** 10.1101/196428

**Authors:** Renaud Schuck, Mary Ann Go, Stefania Garasto, Stephanie Reynolds, Pier Luigi Dragotti, Simon R. Schultz

## Abstract

Multi-photon laser scanning microscopy provides a powerful tool for monitoring the spatiotemporal dynamics of neural circuit activity. It is, however, intrinsically a point scanning technique. Standard raster scanning enables imaging at subcellular resolution; however, acquisition rates are limited by the size of the field of view to be scanned. Recently developed scanning strategies such as Travelling Salesman Scanning (TSS) have been developed to maximize cellular sampling rate by scanning only regions of interest in the field of view corresponding to locations of interest such as somata. However, such strategies are not optimized for the mechanical properties of galvanometric scanners. We describe here the Adaptive Spiral Scanning (SSA) algorithm, which fits a set of near-circular trajectories to the cellular distribution to avoid inertial drifts of galvanometer position. We compare its performance to raster scanning and TSS in terms of cellular sampling frequency and signal-to-noise ratio (SNR). Using surrogate neuron spatial position data, we show that SSA acquisition rates are an order of magnitude higher than those for raster scanning and generally exceed those achieved by TSS for neural densities comparable with those found in the cortex. We show that this result also holds true for *in vitro* hippocampal mouse brain slices bath loaded with the synthetic calcium dye Cal-520 AM. The ability of TSS to ”park” the laser on each neuron along the scanning trajectory, however, enables higher SNR than SSA when all targets are precisely scanned. Raster scanning has the highest SNR but at a substantial cost in number of cells scanned. To understand the impact of sampling rate and SNR on functional calcium imaging, we used the Cramér-Rao Bound on evoked calcium traces recorded simultaneously with electrophysiology traces to calculate the lower bound estimate of the spike timing occurrence. The results show that TSS and SSA achieve comparable accuracy in spike time estimates compared to raster scanning despite their lower SNR. SSA is an easily implementable way for standard multi-photon laser scanning systems to gain temporal precision in the detection of action potentials while scanning hundreds of active cells.

## 1. INTRODUCTION

There has been extensive development of new optical neuroimaging techniques in the last two decades. Since the emergence of multi-photon laser scanning microscopy (MPLSM) in the 1990s [1, 2], synthetic and genetically encoded calcium fluorescent indicators [3, 4] have been used *in vitro* and *in vivo* to monitor the activity of 10s to 1000s of neurons at subcellular resolution [5–7], typically by galvanometric scanning of a point focus throughout the tissue. However, the point scanning nature of this technique limits temporal resolution, and this becomes more pronounced with an increasing number of scanned cells. An important goal of large-scale two photon calcium imaging is to extract time-series signals from as many cells as possible, with sufficient signal to noise ratio (SNR) to detect action-potential induced calcium transients, and sufficient sampling rate to detect calcium transient onset time accurately. There is thus, the need to develop methods which maximise the cell count, sampling rate, and SNR, subject to the trade-offs between these quantities.

The bandwidth limitations of galvanometric scanning systems have motivated the development of inertialess scanning microscopes based on acousto-optic deflectors, and scanless microscopy systems [8]. However, such technology adds considerable complexity to a microscope, and cannot easily be retrofitted into legacy multiphoton microscopes. Resonant scanning galvos allow for increased sampling rates. However, as they still spend much time scanning non-interesting regions of tissue, SNR, which depends on the number of photons collected from structures of interest, suffers. Here, we study the performance of scanning approaches that allow sampling of targeted regions of interest (ROIs) and which can be used with standard, commercial two photon microscope hardware, provided that the microscope software provides the capability to drive the galvos along a pre-defined trajectory.

To increase temporal resolution in galvanometric MPLSM, one strategy is to direct the focus from one target cell to the next along a shorter targeted scanning path, avoiding sampling uninteresting regions of tissue. One implementation of such a strategy, Targeted Path Scanning (TPS) [9], was used to achieve sampling rates of 100 Hz with single action potential (AP) sensitivity, while scanning individual neurons sparsely distributed over the entire hippocampus of a juvenile rat (*>* 1.5 mm). More recently, Sadovsky *et al.* [10] introduced the Heuristically Optimal Path Scanning (HOPS) algorithm. In this approach, the Travelling Salesperson Problem is solved to find the shortest path between all ROIs. In this paper, we refer to Travelling Salesperson Scanning (TSS) algorithms generically (with HOPS being the specific implementation in [10]). With TSS, the fraction of time spent scanning ROIs can be increased from around 4% to 40% when compared to raster scanning, achieving sampling frequencies of 150 Hz for a population of 50 neurons in the somatosensory cortex [10].

However, as galvanometric mirrors are inertial systems, they cannot be driven above a certain speed; otherwise, they fail when scanning sharp turns in the trajectory. Motivated by this, we have developed the Adaptive Spiral Scanning (SSA) algorithm, first reported in [11], and described in more detail here. SSA is based on a simple galvo model, and fits a set of near-circular trajectories to the somatic locations of a population of neurons. In [11], we demonstrated by simulation that high sampling rates can be achieved with minimal-inertia trajectories for typical neural densities found in the visual cortex, somatosensory cortex and hippocampus. In this paper, we present a full description of the implementation of this system, validate the approach by simultaneously performing whole cell patch clamp electrophysiology with two photon scanning of hippocampal and cortical brain slices, and compare its performance with TSS, as well as raster scanning.

To compare signal quality between different scanning strategies, we employ a new (in this context) procedure: calculating the Cramer-Rao Bound (CRB) [12, 13] of the uncertainty in the AP onset time, given the sampling rate (*f*_*s*_) and SNR of the analyzed time series, and the electrophysiological recorded ”ground truth” signal. This metric allows us to demonstrate the effect of varying *f*_*s*_ and SNR, through manipulations of the two photon scanning path employed, on precisely transcribing the activity of large neural networks.

## 2. MATERIALS AND METHODS

### 2.1. Path generation for targeted scanning strategies

Scan paths were generated in MATLAB (Mathworks Ltd), and tested with a standard MPLSM (SliceScope, Scientifica Ltd) (i) on purely simulated neuron location distributions, (ii) on real ROI distributions corresponding to neural somas from raster-scanned slices, but applied ”offline”, and (iii) on real brain slices.

For two-dimensional (2D) simulated neurons, cell centers were assigned to a uniform distribution of random positions in a circular field of view (FOV) of 300 *μ*m diameter with the condition that cells cannot be superimposed. Scan paths were generated with the TSS and SSA algorithms for distributions of 50, 100, 150, 200 and 250 cells (cell radius 7 *μ*m).

For real neuron distributions, we segmented the calcium movies using a semi-automated algorithm based on user input and on the temporal profile of the pixels. First, we apply temporal edge detector and binary thresholding on a spatially down-sampled movie to individual Points Of Interest (POIs). We thenuse the k-means algorithm to classify these POIs as putative neurons or noise based on the similarity between their temporal activity and a stereotypical calcium transient profile. Finally, for each of the POIs labelled as putative neurons, we compute the average time signals and segment the original calcium movie based on the correlation map with such signal, obtaining ROIs. We only keep those ROIs whose shape and size belong to a biologically plausible range. For some of them (depending on how conservatively we set the algorithm parameters), we also ask for feedback from the user before definitively selecting them as neurons. At the end of this procedure, the final result is shown in a graphical user interface in MATLAB where the user can select additional neurons manually. The ROI centers were then fed into the TSS and SSA algorithms to generate the appropriate scanning path.

Scan paths were then uploaded into the Labview-based program *SciScan* (Scientifica Ltd), which controlled the MPLSM, and were tested at varying *f*_*s*_ and target dwell times. The positions of the galvanometric mirrors (Model 8315Kl, Cambridge Technology) were recorded using an electronic position feedback circuit (Micromax 671xx, Cambridge Technology) allowing us to determine the actual path traversed by the laser, and the number of targets hit or missed. MATLAB code for the TSS and SSA algorithms, and details on how to use them in conjunction with *SciScan* are available at http://www.github.com/schultzlab/scanning.

#### 2.1.1. Travelling Salesman Scanning

In our implementation of travelling salesperson scanning, we followed in most respects the HOPS algorithm developed by Sadovsky *et al.* [10]. The TSS algorithm also starts with ordering of scanning locations. To find a near-optimal solution to the shortest path between cells, we used open source genetic algorithm travelling salesperson code, *tsp ga.m*, written by Joseph Kirk, and made available via http://www.mathworks.com/matlabcentral. This code computes a near optimal solution to the shortest path between all the locations to be scanned, with the condition that each location is only scanned once. After the locations to be scanned have been ordered, we applied current steps to both GSs and sampled the trajectory by integrating the ODE characterizing the mechanical behavior of the GSs. Finally, in the TSS computer path generation, we added a condition that enables the user to choose a fixed number of samples per visited ROI. This allows the galvos to be driven as fast as possible from one cell to another while enabling the user to adjust the dwell time over the target depending on the type of dye used, the inertia of the path and the required SNR for the experiment. The user has, however, to bear in mind that GSs are inertial, inducing a slight delay towards reaching their imposed location. It is therefore necessary to choose a number of samples that allows all the ROIs-even those located on sharp turns - to be reached and sampled with a sufficient amount of time. One TSS scanning cycle is achieved when all the cell locations have been successfully scanned.

#### 2.1.2. Adaptive Spiral Scanning

The SSA algorithm is implemented in MATLAB, as part of a simulation platform for MPLSM scanning algorithms [11]. The algorithm inputs are a set of cell soma locations. The first step of the SSA algorithm consists of sorting cell locations by ascending radius *r*_*i*_ from the center of the FOV. Subsequently, the path generation of the SSA algorithm is based on this ordering of cell locations, and on the numerical integration of a linear, second order, Ordinary Differential Equation (ODE) that approximates the mechanical behavior of galvanometric scanners (GS) [11]. In our linear GS model, the driving force is a current that rotates the rotor and therefore, controls the precise angle of the GS mirror. Consequently, driving two GSs with two sine waves results in driving the focal point of the MPLSM in a circle, minimizing the impact of the inertia on the scanning system. The SSA algorithm is initiated by integrating the ODE with two sine waves applied to both GSs with an amplitude corresponding to the radius *r*_1_ of the first cell to be scanned. The radius of the scanned circle is then updated after the first cell location is visited with a number of samples defined by the user. After every cell location visit, the radius index is incremented by a scalar *j* satisfying the equation *r*_*k*_ = *r*_*i*_ + *r*_*j*_ where *k* is the new radius index, *i* the current radius index of the scanned cell and *j* the number of neurons scanned (with a sufficient number of samples) while scanning circle *i*. If *k* = *N* where *N* is the number of cell locations and all the cells have been scanned, then *r*_*k*_ = *r*_1_ and a new cycle starts.

### 2.2. In-vitro cell loading of hippocampal slices with calcium indicator dyes

All animal experiments were performed under institutional guidelines and were approved by the Home Office (UK) and were in accordance with the UK Animals (Scientific Procedures) Act 1986 and associated guidelines. Juvenile wild-type mice (C57Bl/6 P13-P21) were anaesthetized with isoflurane. Depth of anaesthesia was assessed by loss of pedal withdrawal reflex prior to decapitation procedure. Brain slices (400 *μ*m thick) were horizontally cut in 1-4°C oxygenated (95% O2, 5% CO2) slicing artificial cerebro-spinal fluid (sACSF, containing in mM: 0.5 CaCl_2_, 3.0 KCl, 26 NaHCO_3_, 1 NaH_2_PO_4_, 3.5 MgSO_4_, 123 sucrose, 10 D-glucose) using a ceramic blade (Camden instrument 7550/1/C), with the vibratome (7000smz, Camden Instrument) adjusted to the following parameters: Δ*z <* 3*μ*m, horizontal slicing frequency 80 Hz and slicing speed 0.06 mm/s. Slices containing the trisynaptic loop of the hippocampus were taken and allowed to rest in oxygenated recovery ACSF (rACSF, containing in mM: 2 CaCl_2_, 123 NaCl, 3.0 KCl, 26 NaHCO_3_, 1 NaH_2_PO_4_, 2 MgSO_4_, 10 D-glucose) for 30 min at 37°C. Subsequently, the hippocampal slices were incubated and ”painted” for 30 min at 37°C in a dark oxygenated chamber containing the following solution: 2.5 mL rACSF, 50 *μ*g Cal-520 AM (AAT Bioquest), 2 *μ*L Pluronic-F127 20% in DMSO (Life Technologies) and 48 *μ*L DMSO (Sigma Aldrich). Slices where then washed in rACSF at room temperature for 30 min before being transferred to the recording chamber.

### 2.3. Two-photon imaging and electrophysiology

Imaging was performed using a standard commercial GS based two-photon microscope (SliceScope, Scientifica Ltd) coupled to a mode-locked Mai Tai HP Ti:S laser system (Spectra-Physics) operating at 810 nm with pulse width of 100 fs at 80 MHz. A 40*×*, 0.8 NA water immersion objective was used to locate the dentate granule cells (DGCs) or the nearby entorhinal cortex under oblique illumination. Individual cells were filled through the patch pipette (*R* = 3 to 6 MΩ) with intracellular solution containing in mM: 130 K-gluconate, 7 KCl, 10 HEPES, 4 ATP-Mg, 0.3 GTP-Na, 10 phosphocreatine-Na, 0.1 Cal-520 potassium salt (AAT Bioquest) and 0.1 Alexa594 (Molecular Probes). We allowed the calcium dye to diffuse into the cell for 10-30 min, after which we evoked APs in the patched cells by applying current steps while simultaneously recording membrane potential and obtaining two-photon calcium images. Whole-cell current clamp recordings were performed with a MultiClamp 700B using WinWCP 5.2.0 (University of Strathclyde) while two-photon images were acquired using the Labview based software *Sciscan* associated with the commercial microscope. Imaging power at the specimen did not exceed 15 mW.

### 2.4. Assessing the quality of scanned signals

Having presented three different scanning algorithms: raster scanning, TSS and SSA, each of which can be run at different sampling rates, subject to overall physical limitations, it is essential to compare the quality of the signals sampled. There are two aspects to signal quality that we considered. The first is the overall SNR defined as *A/σ*, where *A* is the peak amplitude of a single calcium transient and *σ* is the standard deviation of the noise. The standard deviation of the noise of each trace was estimated as the sample variance in a region of length 5 s (2100 samples) in which no spikes were detected. We estimated the peak amplitude as the value of *A* that minimized the mean square error between acquired time series with the MPLSM and the times series resynthesized from the pulse model using known AP times from the electrophysiological signal. However, the SNR does not uniquely determine the usefulness of the signals produced; for instance, if one wants to precisely determine the time of occurrence of a set of events (such as calcium transient onsets), it is advantageous to have high *f*_*s*_, whereas a lower *f*_*s*_ might be tolerated if all that is needed is to detect the presence or absence of an event. We analyzed the trade-off between SNR and *f*_*s*_ by taking a second, complementary measure of signal quality: the CRB of the mean squared error in estimating the time of occurrence of an action potential from fluorescence calcium imaging data. This bound gives us the best case scenario for uncertainty in spike time estimation for a given SNR and *f*_*s*_. We calculated the CRB as

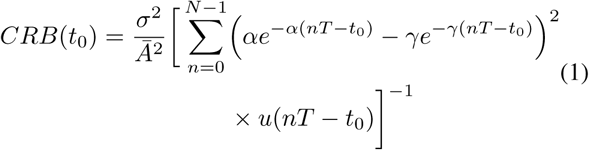

where *t*_0_ is the spike time, determined by whole cell recordings; *σ* is the standard deviation of the noise; *N* indicates the number of samples in the time series, and *T* = 1*/f*_*s*_. *u* is the indicator function where *u*(*nT-t*_0_) = 0 for *nT - t*_0_ < 0 and *u*(*nT - t*_0_) = 1 for *nT - t*_0_ ≥ 0, Ā is the normalized pulse amplitude and *α*, *γ* represent the typical rise and decay parameters of a single calcium transient, respectively. Parameters *α* = 3.18 s^*-*1^, *γ* = 34.49 s^*-*1^ were estimated from the rise and decay time in [4]. Further details of the CRB derivation are provided in the Appendix.

## 3. RESULTS

Both high temporal resolution and SNR are important in order to faithfully transcribe spike trains from the calcium traces of recorded neurons. An important aim of our experiments was, thus, to compare these criteria for raster scanning, TSS and SSA applied to two dimensional scanning for *in vitro* slice experiments.

### 3.1. Sampling frequency dependence on scanning strategy

Calcium transients can be detected when the laser is focused on a neuron while its internal calcium is elevated following an AP. In our *in vitro* experiments we used the synthetic calcium dye Cal-520, which has a rise time of *<* 70 ms and a decay time constant of *<* 800 ms for a single AP [4]. In an FOV of 300 × 300 *μ*m^2^, the sampling frequency achieved by raster scanning is about 3 Hz. Although higher temporal resolution can be achieved, it comes with a substantial reduction in the FOV, and thus, the number of cells that can be sampled. In contrast, targeted 2D line-scanning strategies such as TSS and SSA allow calcium transient detection from a much larger FOV at the same sampling frequency (and for the same FOV, allow higher sampling frequencies and better signal reconstruction). To compare the sampling frequencies of these scanning strategies under well-controlled experimental conditions, we simulated the somatic locations of a random population of cells (drawn from a spatial distribution approximately the same as that of layer 2/3 of mouse primary visual cortex [7]), and scanned these locations with the two photon microscope (Fig. 1a-d).

**Figure 1:**
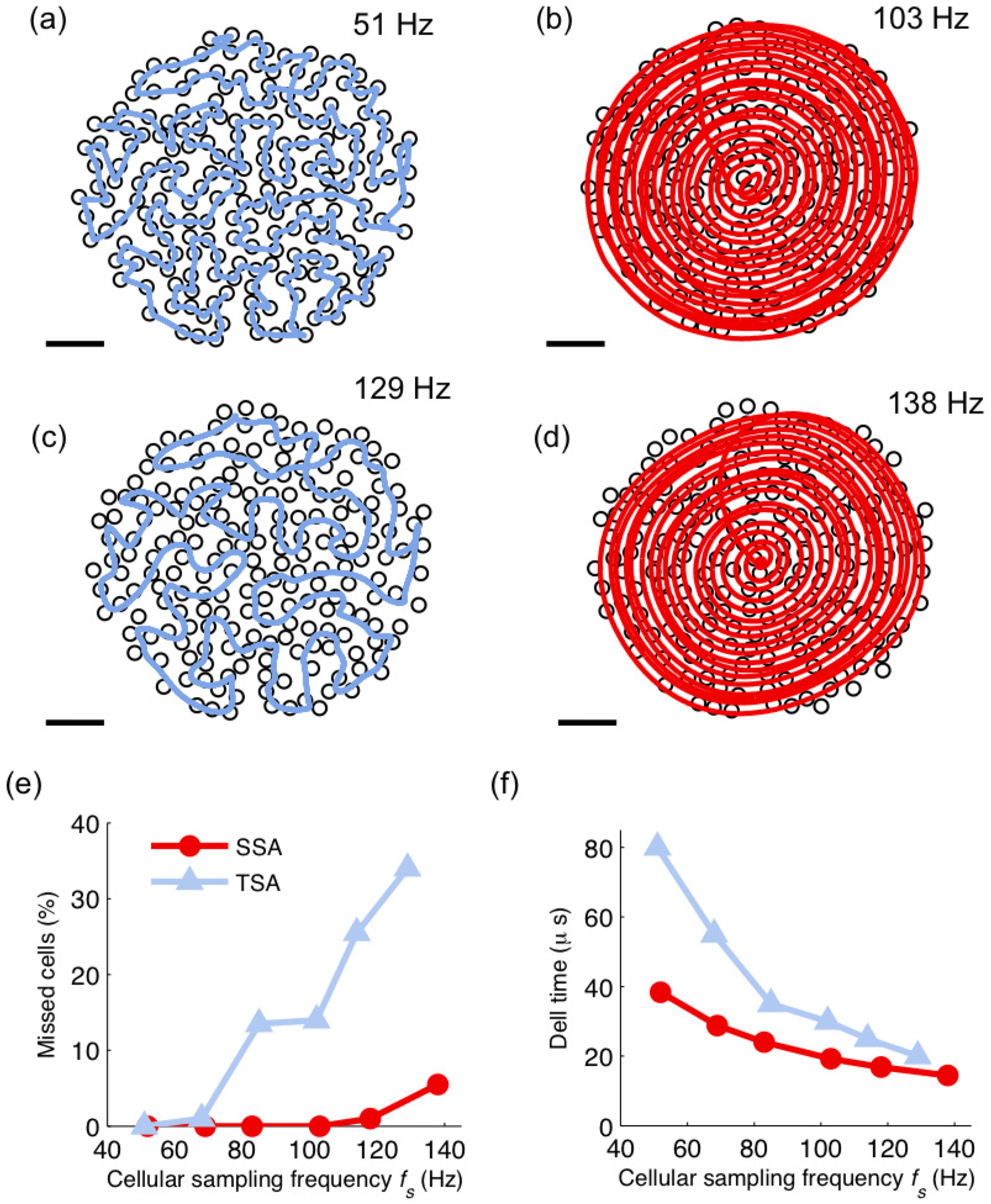
Spiral scanning yields lower cellular miss rates, but travelling salesperson scanning higher SNRs, for the same somatic locations. (a) Recorded galvo positions for one full path of the TSS for 200 neurons at *f*_*s*_ = 51 Hz (all cells are scanned). (b) SSA path for the same neural population in *a*, but with a scan rate of 103 Hz, the maximum for which all targets were visited. (c) TSS scanpath for the same population, with neuron sampling rate increased to 129 Hz. At this rate, 65 cells were missed. (d) SSA scanpath at 138 Hz, 11 targets were missed (scale bar 50 *μ*m for *(a)*, *(b)*, *(c)* and *(d)*. (e) The fraction of target cells missed for increasing *fs* with the same soma locations as above. (f) The average time spent collecting photons from each ROI, for the SSA and TSS paths as a function of increasing *f*_*s*_. Increasing value on the y-axis indicates greater SNR in the resulting time series signals.

Employing TSS on a simulated population of 200 cells in a 300 × 300 *μ*m^2^ area showed that for this (typical) instance of random cell positions, the highest cellular sampling rate *f*_*s*_ allowing the galvos to follow the command voltage and hit all desired targets was 51 *±* 7.4 Hz (Fig. 1a). Any increase of *f*_*s*_ above this value resulted in missed cells, due to the system not being able to follow precisely the desired trajectory (Fig. 1b). Performing a similar assessment on the same neuronal locations for SSA shows that all the targets can be hit at sampling rates up to 103 *±* 6.7 Hz, almost double the maximum *f*_*s*_ for TSS (Fig. 1c). In general, the number of missed cells is lower for SSA compared to TSS (Fig. 1e). However, at any given sampling frequency *f*_*s*_, the microscope spends more time collecting photons from ROIs per sample under TSS (Fig. 1e). This quantity is essentially a measure of the SNR for the resulting time series, and thus, it appears that TSS can be advantageous when maximising sampling rate is not the most important criterion. In the experiment above, the neural density was 2200 neurons/mm^2^. In general, higher sampling rates for SSA (in comparison to TSS) were achieved for target densities exceeding 1400 neurons/mm^2^ (Fig. 2a-d). Automated cell contour detection in mouse somatosensory and visual (V1) cortex from *in vitro* MPLSM high resolution images shows neural densities around 3100 cells/mm^2^ and 2100 cells/mm^2^, respectively [7, 10]. This suggests that when dense sampling of neurons is required, higher sampling rates are achievable using SSA. It should be noted that in some cases, labelling may be sparse (eg. due to use of a cell-type selective promoter), so this provides an upper limit on target density. The dwell time advantage of TSS can be seen to apply regardless of neuronal density (Fig. 2e).

**Figure 2:**
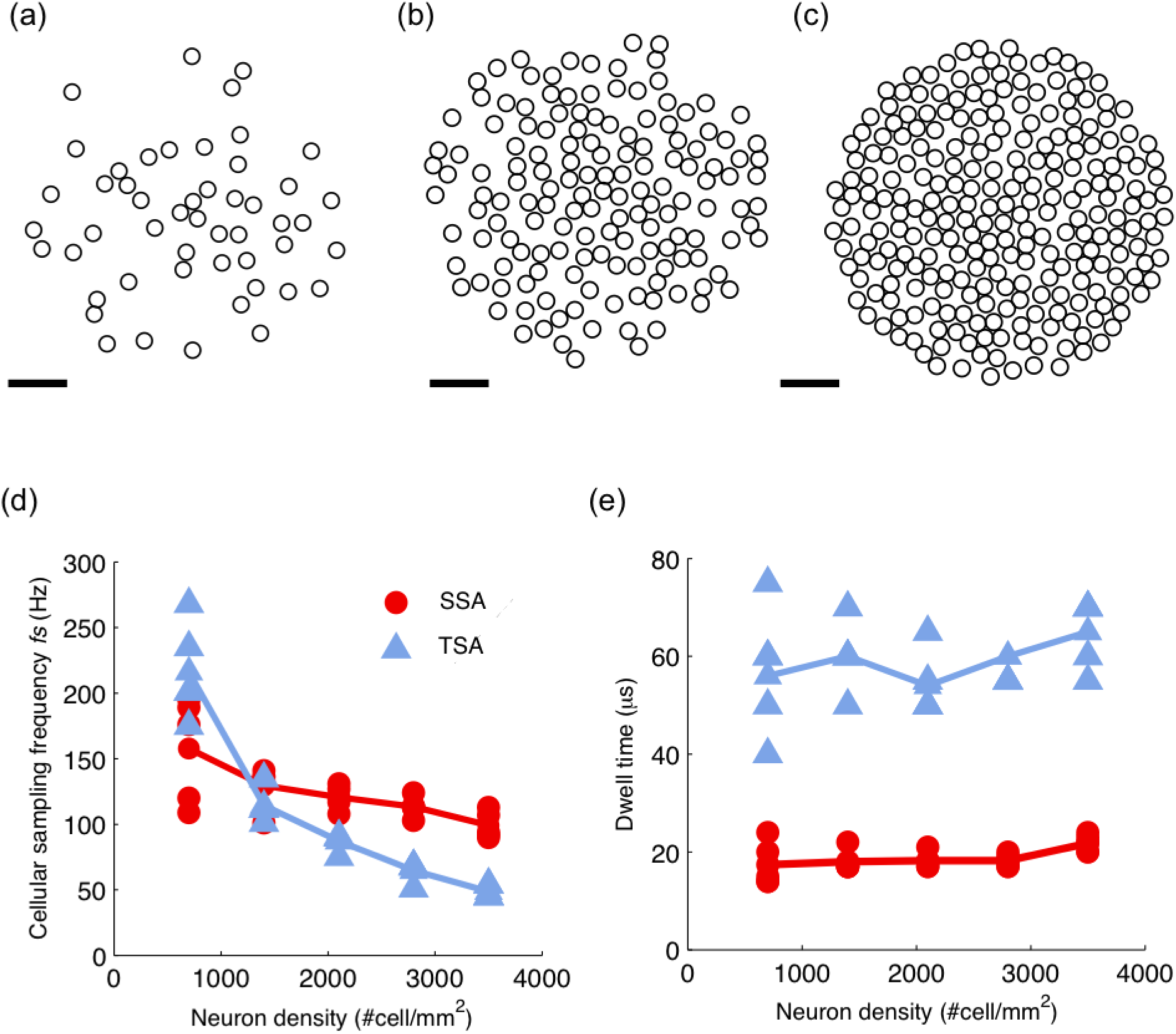
Sampling frequency and dwell time achievable for the TSS and SSA algorithms. (a)-(c) Exemplar spatial distributions of cells (uniformly distributed within a circle of diameter 300 *µ*m), for a total cell count of 50 (*a*), 150 (*b*) and 250 (*c*) cells. These examples correspond to densities of 700, 2100 and 3500 cells/mm^2^, respectively (scale bar 50 *µ*m for *(a)*, *(b)* and *(c)*). (d) The maximum sampling frequency at which all the targets are hit, for TSS and SSA across different neuronal densities (each cell density was tested on five different geometric cell arrangements). (e) Average dwell time over the target across the same neural distributions as used in *(d)*.

To further validate and characterise the performance of the algorithms, we performed *in vitro* two photon imaging experiments in mouse hippocampal brain slices bath-loaded with the calcium dye Cal-520 AM, as described in Methods. Figure 3a shows the dentate gyrus of such a bath-loaded slice. This area, which is densely packed with granule cells, provides an excellent testbed for the platform. We used an initial raster-scan, together with semi-automated cell body localisation (see Methods), to locate individual granule cells that were either spontaneously active or electrically driven during the raster-scan (cyan discs in Fig. 3b). In this particular image, 137 cells were extracted within a FOV of 180 × 180 *μ*m^2^ (corresponding to a density of *∼*4200 cells/mm^2^). TSS allowed scan rates (inverse of time to cover one full cycle back to the same location) of up to 88 Hz before the path started missing ROIs (Fig. 3b shows the path at 88 Hz) Applying the SSA algorithm, we generated a spiral trajectory which passed through all cell bodies at sampling rates up to 117 Hz (Fig. 3b shows the 117 Hz path). For lower cell densities, however, we found that often TSS would reach higher sampling rates than SSA before missing cells. For instance, at a density of 50 cells per 180 × 180*μ*m^2^ (1500 cells/mm^2^), SSA was able to reach 165 Hz, whereas TSS was able to reach 208 Hz (Fig. 3d). Both scanning algorithms yield high quality calcium transient recordings (Fig. 3e, f).

**Figure 3:**
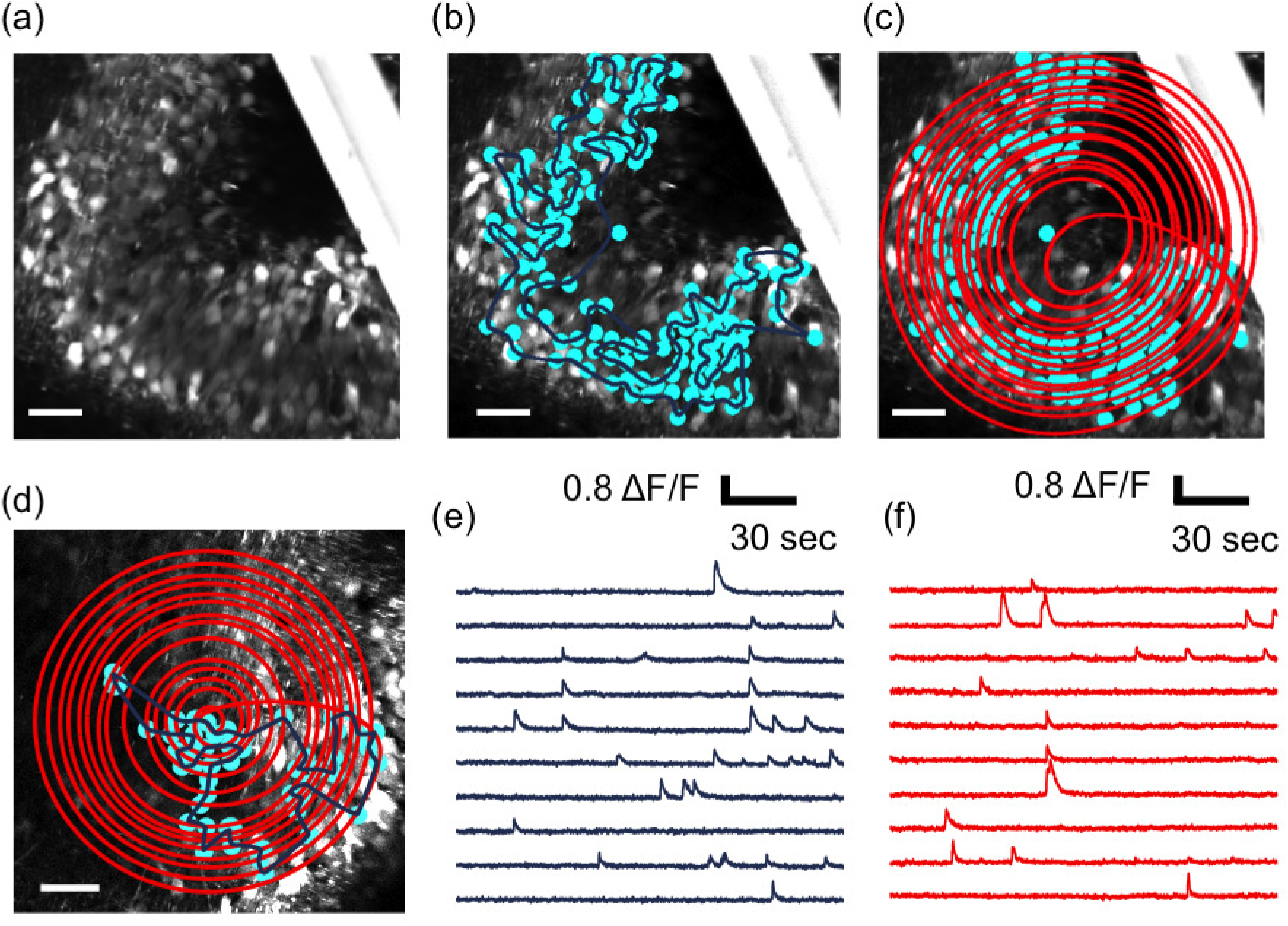
Scanning patterns for fast acquisition of calcium signals. (a) Raster-scanned image of dentate granule cells used to locate targets for scanning. (b) 137 ROIs (in cyan) are shown overlaid, indicating granule cells that were spontaneously active or electrically activated during raster-scanning for target acquisition. The overlaid dark blue trace indicates the recorded galvo trajectory for the highest *f*_*s*_ (88 Hz) at which TSS is able to hit all targets. (c) Overlaid SSA path (red), at maximum *f*_*s*_ (117 Hz) to hit all targets. (d) Frame scanned image of dentate granule cells with 50 ROIs (in cyan) overlaid with scanning paths for maximum *f*_*s*_ for TSS (208 Hz, dark blue) and SSA path (165 Hz, red), scale bar 25 *µ*m for *(a)*, *(b)*, *(c)* and *(d)*. (e) Spontaneous activity acquired from 10 (of 50) granule cells with TSS at 208 Hz (traces 40-point median filtered for visualization only). (f) Spontaneous activity traces from the same granule cells acquired with SSA at 165 Hz (traces 50 point median filtered to match temporal regularisation in *(e)*).

Missed targets can be seen at sharp turn locations in the TSS trajectory, indicating that the inertia of the galvanometric scanners do not allow them to sharply change direction above a certain speed. To examine this further, we segmented 50 ROIs corresponding to spontaneously active dentate granule cells in 180 × 180*μ*m^2^ and scanned them with TSS at *f*_*s*_ rates ranging from 166 Hz to 885 Hz (Fig. 4a,b). Consider the scanned granule cell encircled in orange in Figure 4b: the target ROI is just grazed for *f*_*s*_ = 425 Hz, and is missed entirely above this scanning frequency. The recorded time series for this example is shown in Figure 4c (no median filtering). At 885 Hz, no functional signal was visible, as the scanpath did not intersect with the ROI. More of the functional signal is retrieved at 425 Hz and 166 Hz when 36 *μ*s and 122 *μ*s, respectively, were spent collecting photons per cycle (dwell time). Note that in TSS the dwell time can be artificially increased by causing the focus to stop for a period in the centre of the cell. This, however, comes with a substantial penalty in terms of sampling rate.

**Figure 4:**
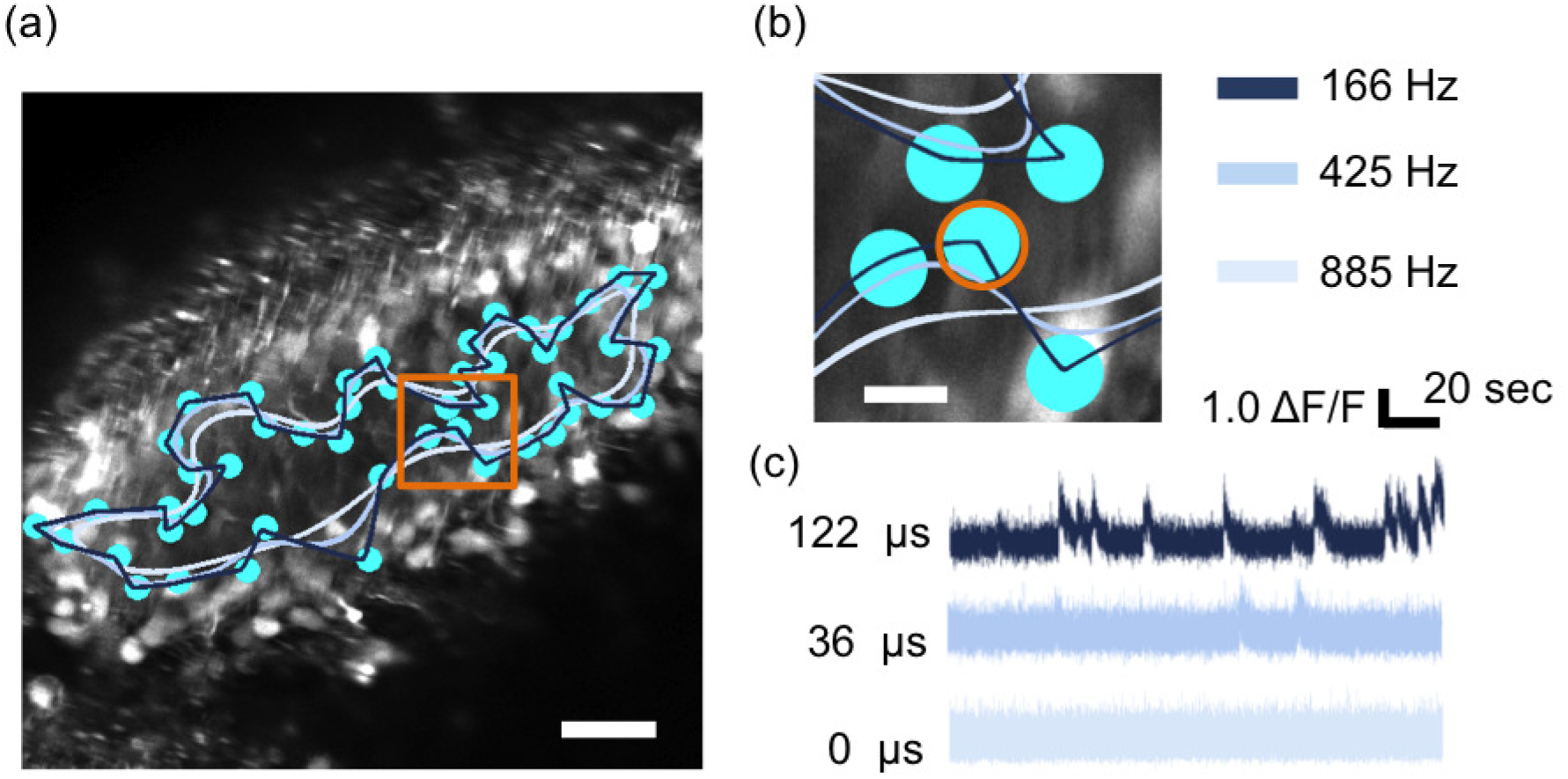
High sampling rates in TSS lead to ROI dropping. (a) Raster scan of an imaged region of dentate gyrus. Spontaneously active neurons are indicated by cyan discs. The 3 different TSS paths shown indicate the trajectory followed by the MPLSM focal point when the galvos were driven at increasing speed (scale bar 25 *µ*m). (b) Expansion of the orange square in (*a*); path colour indicates sampling rate (scale bar 10 *µ*m). (c) Time-series of the orange-highlighted neuron for different dwell times corresponding to the different scanning frequencies in (*b*).

### 3.2. Signal quality

#### 3.2.1. SNR dependence on scanning strategy

We saw in section 3.1 that while the SSA algorithm has (depending on the exact distribution of cells) some advantages for sampling rate, TSS tends to produce higher dwell times over each neuron for the same neuron density. This dwell time could be further increased by ”parking” the laser over a cell for longer. However, this comes at the expense of sampling rate. The duration of time spent collecting photons from each target structure provides an indirect way to compare the SNR (all other things being equal). SNR is critical to the fidelity with which APs can be reconstructed from calcium transient time series. Therefore, we analysed the extent to which it can be improved by increasing the dwell time over each neuron in TSS.

To examine this, we used the TSS paths at different sampling frequencies for functional calcium imaging of cortical cells while simultaneously doing whole cell patch-clamp electrophysiology on one of the cells in the scanning path (Fig. 5a). This allowed us to elicit individual APs from the patched cell, in current clamp mode, in order to measure SNR. Increasing numbers of APs were elicited, yielding calcium transients of increasing amplitude, and correspondingly, SNR (Fig. 5b). For a relatively small number of APs (up to 8), signal amplitude increased approximately linearly with the number of APs elicited for the three TSS paths (Fig. 5b). We measured the SNR per spike by taking a linear fit of the plot of SNR (calculated as described in Methods) versus the number of APs elicited (Fig. 5c). The slope of the linear fit can then be interpreted as the SNR per spike, that is, a measure of the signal quality independent of the number of APs in a given event. The increase of the slope values in Figure 5c between the three TSS paths clearly shows that the SNR per spike increases with dwell time (time spent collecting photon from the ROI) albeit at the expense of sampling frequency.

**Figure 5:**
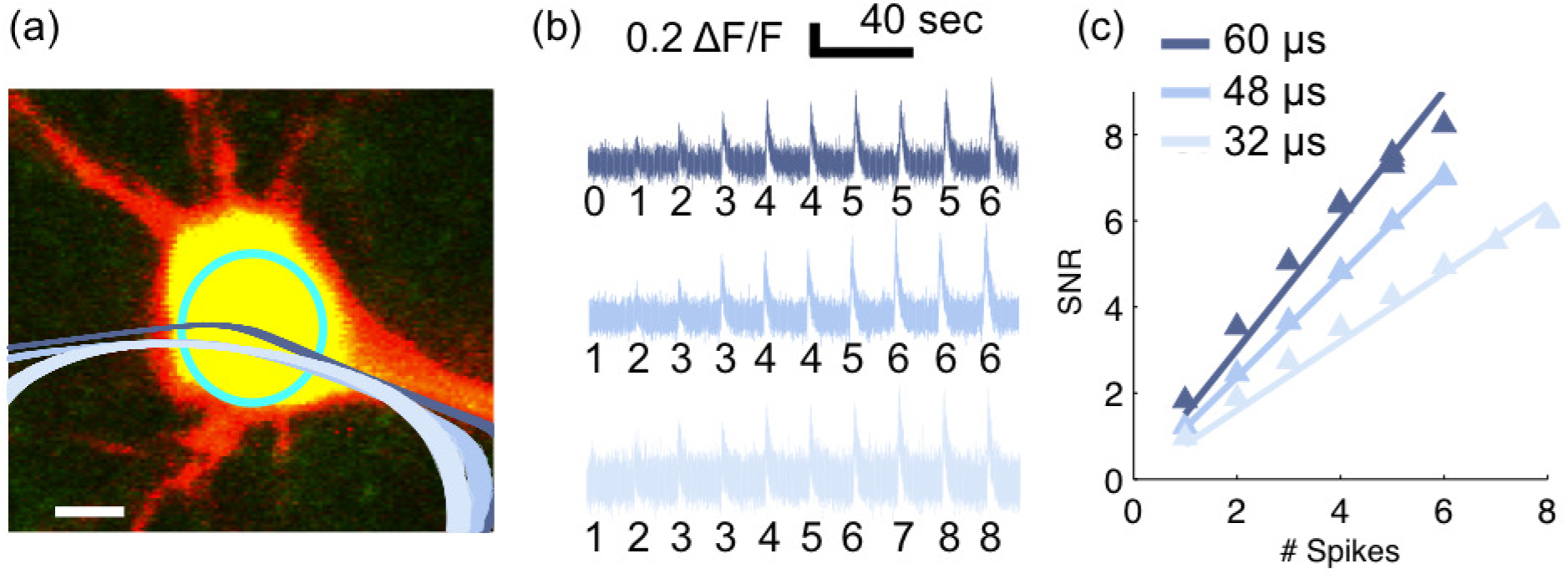
Photon collection with the TSA. (a) Zoom in on a whole cell patch of a cortical pyramidal cell with the associated ROI (cyan circle) and three TSS paths overlaid: 102 Hz (dark blue), 137 Hz (medium blue) and 208 Hz (light blue) (scale bar 10 *µ*m). (b) Calcium time series associated to the TSS paths in (*a*) and their corresponding number of APs. (c) SNR evolution with an increasing number of APs for the three TSS paths shown in (*a*) and corresponding to the calcium traces shown in (*b*). The photon collection time (dwell time) from the patched cell was: 60 *µ*s (dark blue data points), 48 *µ*s (medium blue data points) and 32 *µ*s (light blue data points). For each TSS path, a linear least square fit has been used on the SNR values.

To further compare the trade-off between SNR and sampling frequencies amongst raster scanning, TSS and SSA, we used the same technique of simultaneous functional calcium imaging and whole cell patch clamp recording of one cell on the scanning path. For each patched cell (*n* = 4 in our study) we could define multiple cell geometries (with different cell counts) and therefore, try out multiple scanning paths within a 300 × 300 *µ*m^2^ FOV. To determine and visualize the scanning paths of TSS and SSA, we overlaid them over the recorded tissue. If at the depth of the studied patched cell, surrounding neurons were not well loaded with the Cal-520 AM bath loading technique, we chose random ROIs to be scanned by TSS and SSA. We found our Cal-520 AM bath loading technique most effective down to *∼* 30 *µ*m under the slice surface for mice *<* PND21 (see Methods). Below this depth, the loading of the tissues decreases down to –50 *µ*m where it stops. We filled the patched neuron with Cal-520 potassium salt (see Method) to ensure that the cell was well-loaded.

Figure 6a shows a representative two-photon image of a cortical pyramidal cell while Figure 6b shows the overlaid TSS and SSA scanning paths. The recorded somatic potential and the corresponding calcium trace from raster scanning at 10 Hz are shown in Figure 6c. This scanning frequency was achieved by raster scanning over a much smaller FOV, 180 x 180 *µ*m^2^. The calcium traces from TSS and SSA scanning over the entire FOV are shown in figure 6d. For all the scanning strategies, we also found an almost linear increase in the SNR with the number of APs (up to 8). This is shown in a linear fit of the plot of SNR versus number of APs for three examples, with different scanning algorithms and sampling rates, in Figure 6e. Note that these are examples taken at different sampling rates, and should not be compared directly. To appreciate the trade-off between SNR and sampling rate for the three scanning algorithms, we then plotted the cumulative data for SNR per spike versus sampling rate for the different experiments (Fig. 6f). The three examples from Figure 6e are indicated in Figure 6f by orange symbols. Overall, it is apparent that raster scanning typically achieves the highest SNR per spike: 1.49 (median value amongst *n* = 14 trials), importantly, at a substantial cost in sampling frequency and FOV (or number of cells sampled). In these examples, TSS and SSA scanning achieved similar SNR per spike of 0.75 (median of *n* = 9 TSS paths) and 0.74 (median of *n* = 14 SSA paths), respectively. The similarity in SNR for TSS and SSA in these examples is due to the TSS path being driven at high sampling rates and therefore reducing the dwell time spent on the patched neuron. Path precision analyses on galvo positions also show that most of the high sampling frequencies achieved with the TSS paths are missing ROIs (Fig. 6f). For the same neural distributions, the SSA paths were driven at relatively conservative *f*_*s*_ and therefore did not miss any ROIs (Fig. 6f).

**Figure 6:**
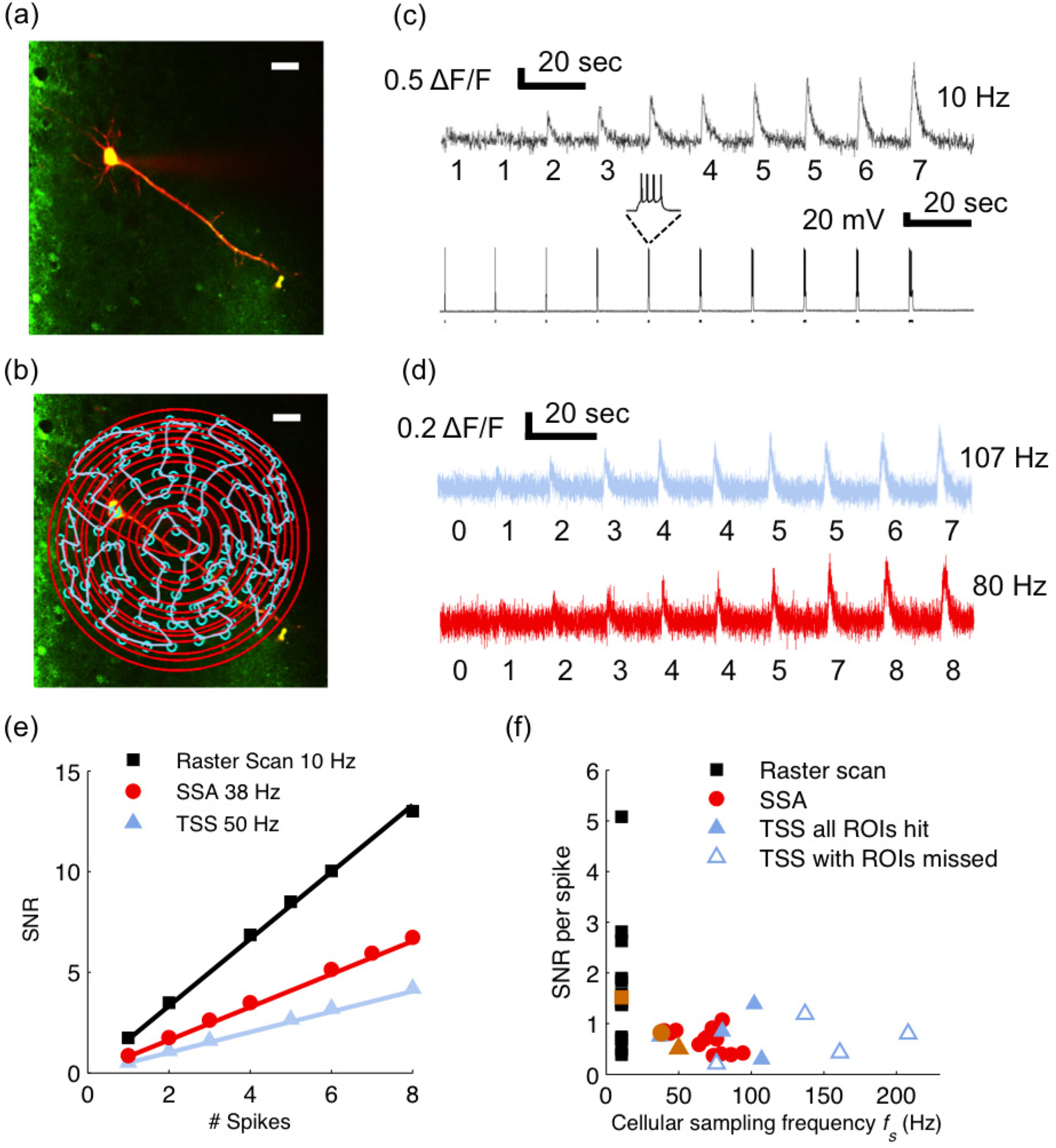
Simultaneous functional calcium imaging and ground truth electrophysiology recordings for raster scanning and scanning strategies. (a) High resolution image showing a wholecell patch of a cortical pyramidal cell (scale bar 25 *µ*m). (b) High resolution image showing the SSA (in red) and TSS (in light blue) scanning trajectories for 129 neurons. The locations of the neurons in cyan (except the cell that is patched) have been chosen arbitrarily as the neurons were not well loaded with Cal-520 AM) at this imaging depth (scale bar 25 *µ*m). (c) Representative example of recorded somatic potential (bottom) and raster scanning associated time series (top) sampled at 10 Hz. Black bars under recorded somatic potential indicate the current step duration in the stimulation protocol. The number of APs evoked is written under each calcium transient. (d) Representative example of times series sampled at 107 Hz for TSS (in light blue) and 80 Hz for SSA (in red). The numbers of APs at every stimulation is written under each trace. (e) Example of SNR evolution with an increasing number of APs for raster scanning, TSS and SSA. For each type of scan, a linear least square fit has been used on the SNR values. (f) SNR per spike values for multiple (n=14) raster scans at 10 Hz and multiple SSA (n=14) and TSS (n=9) scans for cellular sampling frequencies within [38 Hz to 208 Hz]. The filled data point markers represent scanning paths where all the ROIs are hit whereas the open ones represent the paths with missed targets along the scanning trajectory. The orange markers correspond to the example in (*e*).

### 3.2.2. Lower bounds on the uncertainty of AP timing

The quality of signals obtained by multiphoton fluorescence calcium imaging can be measured in various ways. One way is to measure the root mean square (RMS) deviation between the measured fluorescence time series, and the true underlying calcium signal -if one had access to it. Unfortunately, however, this is not the case. In any event, for many applications we are particularly interested in reconstructing spike trains, and thus, in the time at which AP-induced calcium transients occur. Relevant ground truth for this situation can be obtained, by performing whole-cell patch clamp electrophysiology simultaneously with imaging, as described above. The question then is how well the timing of each AP can be reconstructed from the scanning data [14]. One way to assess that would be to apply a particular spike train reconstruction algorithm, and to measure the RMS error in the timing of spikes. However, the results would apply only to that particular reconstruction algorithm. A more general approach, which we take here, is to compute the Cramer-Rao lower bound (CRB) of the variability with which spike timing can be extracted using any algorithm, based upon the measured properties of the recorded signals. Details of this calculation are provided in Methods and Appendix 4. The resulting quantity describes the temporal precision of the scanned signals, and can thus be used to compare different scanning algorithms in this respect.

We analysed the experiments from the previous section in which whole-cell patch clamp electrophysiology was used to evoke accurately timed APs while one of the three scanning algorithms was used to record calcium transients. For each acquisition, we calculated the CRB of the uncertainty in spike time estimates from the time series. Figure 7 shows the interplay between SNR and *f*_*s*_ on the CRB. Figure 7a shows the CRB - SNR plane and we see that for each scanning algorithm the CRB value decreases, that is, the temporal precision of the acquisition increases, when the SNR value increases. For a given uncertainty in AP timing, raster scanning requires the highest SNR among the three. That is, TSS and SSA are able to achieve comparable uncertainties in spike time estimates as raster scanning despite their lower SNR (Fig. 7a). This is due to their high sampling frequencies. Similarly, in the CRB - *f*_*s*_ plane in Figure 7b, we see that raster scanning can achieve AP uncertainty less than 20 ms only at SNRs greater than 3. In contrast, AP timing uncertainty below 20 ms was achievable for SNRs less than 1.5 with both TSS and SSA.

**Figure 7:**
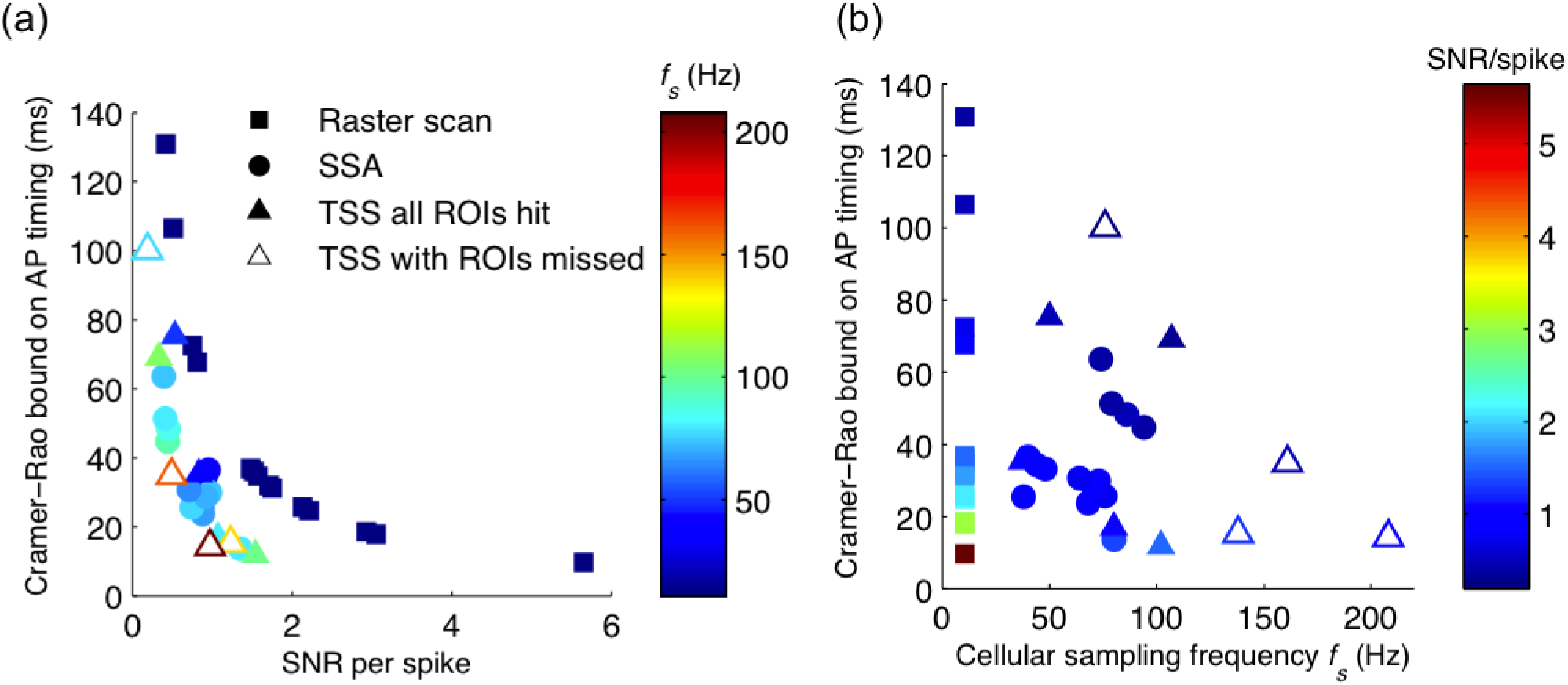
Uncertainty in timing of reconstructed action potentials depends on both SNR and sampling frequency. (a) The Cramer-Rao lower bound on the uncertainty with which the time of an action potential can be estimated, as a function of SNR (abscissa) and sampling frequency (colour scale), for raster scanning (n=14 cells), TSS (n=9 cells) and SSA (n=14 cells). Ground truth spike timing was obtained by simultaneous whole-cell patch clamp electrophysiology. (b) The same dataset visualised with sampling frequency on the abscissa and SNR on the colour scale.

## 4. DISCUSSION

We have described in detail the Adaptive Spiral Scanning algorithm for use with a standard commercial galvo-based multiphoton microscope and have demonstrated its capability for recording high quality, high sampling rate (100s of Hz) time series signals from many cells simultaneously. We have compared its performance in terms of sampling rate and SNR to raster scanning and the Travelling Salesperson Scanning algorithm. We have found that SSA outperforms raster scanning by an order of magnitude in terms of achieved *f*_*s*_ for the same population of neurons. It likewise outperforms TSS in *f*_*s*_ for common neural densities encountered in *in vitro* experiments (visual cortex, somatosensory cortex, GCs of the hippocampus). We have also found the SNR of TSS to be higher than that of SSA as TSS spends, on average, more time per cell collecting photons since it allows the laser to be ”parked” on each ROI. However, the SNR of raster scanning exceeds those of both scanning strategies TSS and SSA as it scans the entire neuron. The low sampling rate for raster scanning is sufficient to capture calcium events. However, the cost for raster scanning is a much smaller FOV and thus, fewer cells that can be sampled. For instance, a 300 × 300 *µ*m^2^ region may be sampled at *∼*3 Hz by raster scanning, whereas the same region may be sampled by SSA at *∼* 100 Hz. For a larger FOV (and hence, a larger neural population), e.g 500 × 500 *µ*m^2^, the raster scanning sampling rate decreases to *∼* 1 Hz. Although this frequency may still allow some calcium events to be captured, there will be considerable loss in the accuracy of spike timing estimates.

We have also compared the different scanning strategies in terms of signal quality of the calcium traces. This can be measured by a number of different metrics [14–17]. In our study, we used the Cramer-Rao Bound to measure the temporal precision between the onsets of the calcium transients and their associated APs emitted by the neuron. This metric shows the weight of the *f*_*s*_ and SNR in the temporal precision of the acquisition (see 2.4 and 4). The values of the CRB represent the lower bound of an AP time estimate from the calcium imaging data. We found that to attain the same temporal precision, raster scanning needs a significantly higher SNR than either TSS or SSA. This is due to the *f*_*s*_ of the SSA and TSA being at least an order of magnitude higher than the *f*_*s*_ of raster scanning for the same number of neurons scanned.

The measure of signal quality and the choice of the scanning type for functional calcium imaging is heavily dependent on the experimental setup. In *in vitro* experiments, it strongly relies on the ability to induce the dye into the tissue with the bath loading technique. The latter depends on the composition of the bath and the age of the animal [18]. In older brain slices (*>*PND 21), protective recovery slicing methods [19], bulk loading [20] or even newly developed bath loading techniques [21] help to induce the dyes into the tissue even if the amount of dye induced is less then in juvenile mice. Therefore, using a longer dwell time per scanned neuron (achievable with the TSS) might be necessary to obtain a good SNR when imaging older dye loaded brain slices. The quality of the signal is also determined by the choice of the dye. Recent development of genetically encoded voltage indicators [22, 23] show major improvements regarding kinetics when compared to calcium indicators. In the case of voltage indicators, fetching a single fluorescent event is only achievable at high sampling rates (achievable with TSS and SSA) as its kinetics is relatively close to the original AP. However, it remains to be seen if scanning strategies enable sufficient SNR to precisely detect a voltage fluorescent event engendered by a single AP. All these experimental aspects highlights the need to develop adaptable scanning strategies for standard MPLSMs to enable the fine tuning of scanning parameters and obtain the best signal quality considering the general experimental framework limitations.

Moreover, another positive aspect resulting from scanning strategies is the minimal coverage of the imaged FOV in comparison to raster scanning: only one line segment passes through each ROI for each full trajectory thus limiting the overall heating of the tissue and bleaching of the dye [24].

In addition to improving the functional fluorescent signal quality acquired with a standard MPLSM of large neural networks, these scanning strategies only require software development and hence are not expensive and easily transferable from one setup to another.

## Appendix: Crameér Rao bound for spike detection from calcium imaging data

We model a fluorescence signal consisting of one spike at time *t*_0_ as

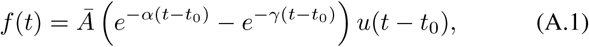

where *u*(·) is the indicator function and the parameters{ Ā*, α, γ* } define the pulse amplitude and the speed of the pulse’s rise and decay. We refer to Ā as the normalized pulse amplitude — the normalization ensures *f* (*t*) has peak value Ā. We have *N* noisy samples 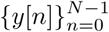, such that

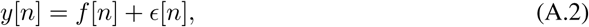

where *∊*[*n*] are samples of a zero-mean Gaussian process with standard deviation *σ*.

The Crameér-Rao lower bound on the variance of an estimate of parameter *t*_0_ from samples *𝒴*[*n*] of the form in equation (A.2) is the Inverse of the Fisher Information *I*(*t*_0_), where

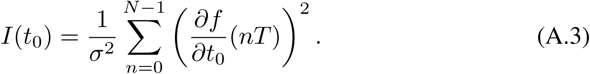

Using the chain rule for differentiation and equation (A.1), we obtain

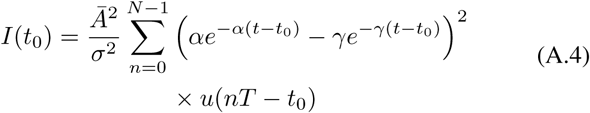

The Cramér Rao bound is thus

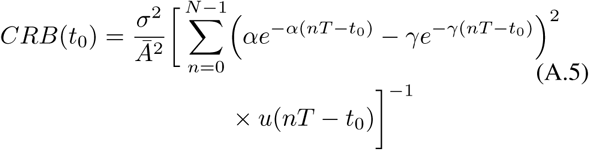

## Acknowledgments

This work was funded by EU Marie Curie Initial Training Network 289146 (”NETT - Neural Engineering Transformative Technologies”), by the UK Biotechnology and Biological Sciences Research Council (award BB/K001817/1), and by a Royal Society Industry Fellowship to SRS.

